# Topology independent structural matching discovers novel templates for protein interfaces

**DOI:** 10.1101/235812

**Authors:** Claudio Mirabello, Björn Wallner

## Abstract

**Motivation:** Protein-protein interactions (PPI) are essential for the function of the cellular machinery. The rapid growth of protein-protein complexes with known 3D structures offers a unique opportunity to study PPI to gain crucial insights into protein function and the causes of many diseases. In particular, it would be extremely useful to compare interaction surfaces of *monomers*, as this would enable the pinpointing of potential interaction surfaces based solely on the monomer structure, without the need to predict the complete complex structure. While there are many structural alignment algorithms for individual proteins, very few have been developed for protein interfaces, and none that can align only the interface residues to other interfaces or surfaces of interacting monomer subunits in a topology independent (non-sequential) manner.

**Results:** We present InterComp, a method for topology and sequence-order independent structural comparisons. The method is general and can be applied to various structural comparison applications. By representing residues as independent points in space rather than as a sequence of residues, InterComp can can be applied to a wide range of problems including: interface-surface comparisons, interface-interface comparisons and even comparisons of small molecule ligands. We demonstrate a use-case by applying InterComp to find similar protein interfaces on the surface of proteins. We show that InterComp pinpoints the correct interface for almost half of the targets (283 of 586) when considering the top 10 hits, and for 24% of the top 1, even when no templates can be found with the already available sequence-order dependent methods like TM-align.

**Availability:** The program is available from: *http://wallnerlab.org/InterComp*

**Supplementary information:** Supplementary data included in the pdf.

## 1 Introduction

Proteins are involved in almost all processes in cells and have evolved to interact with a range of other molecules, such as proteins, DNA, RNA, or small molecules. The study of how proteins interact with these molecules offers important insights into the function of proteins, the way they operate, and possible causes of disease (Alberts, 1998; Jeong *et al*., 2001; Li *et al*., 2004).

Proteins interact with other molecules by making direct physical contact through specific residues on the protein surface. These residues constitute the *interface* of a protein. A variety of interfaces have been experimentally identified and have been found to vary both in shape and residue composition (Davis and Sali, 2005). Interfaces can be stable, as for the multiple chains of the ribosome, or transient, as for many proteins involved in signalling pathways. The same interface can interact with multiple molecules and an interaction can also require multiple interfaces (Bomsztyk *et al*., 2004; Cohen, 2002; Han *et al*., 2004).

To predict how proteins interact with other molecules it is of fundamental importance to know where the interfacial residues are located on their surface. For example, in the case of protein docking, it has been shown that it is relatively easy to dock proteins using *template-based* docking techniques if a similar interaction has been experimentally determined (Tuncbag *et al*., 2012; Kundrotas *et al*., 2012; Zhang *et al*., 2013; Mirabello and Wallner, 2017).

A number of template-based docking methods have been developed in the last few years. Some methods use sequence- or profile-based alignments to match two target protein sequences to the sequences of two protein chains that are part of an experimentally solved quaternary structure (Chen and Skolnick, 2008; Mukherjee and Zhang, 2011). When a match is found, the structure and mutual position of the two protein chains can be used as templates to model the interaction of the targets. Unfortunately, this approach inherits the same drawbacks as template-based modelling of protein monomer structures, and when the pairwise sequence identity drops below 30% it is difficult to obtain a reliable prediction (Aloy *et al*., 2003).

Other methods are based on *structural templates*. The first step in these methods is usually to build the monomer structure of each molecule in the complex separately. Then, the structure of each monomer is aligned to libraries of known complexes, and cases where the two target monomers are structurally similar to monomers of a known complex can be used as a template for the interaction. This improves the coverage of modellable multimers. Since the structure of proteins is more conserved than the sequence, more distantly related homologs can be found using structure (Aloy and Russell, 2002; Zhang *et al*., 2013; Mirabello and Wallner, 2017). However, a drawback of the methods based on structure is that both the target monomers need to be overall structurally similar to their templates. At the same time, it has been shown that an interaction is only specific to the shape and chemical composition of the patches of residues directly involved in the interaction, i.e. the interfacial residues and not the overall structural scaffold. In fact, some proteins interact through the same type of interface while differing substantially in their overall structural similarity (Keskin and Nussinov, 2007).

In addition, the number of different spatial arrangements of residues in protein interfaces seems to be lower than the number of different protein folds, and studies have shown that the space of interface structures is already covered to a large degree in the known structures in the Protein Data Bank (Gao and Skolnick, 2010b; Kundrotas *et al*., 2012). Thus, in principle it should be possible to find template interfaces for most unknown interactions from the structure of single interacting monomers, particularly if the alignment is limited to the interfacial regions of proteins. But aligning only interface residues is not trivial and methods adopt a mixed approach to this problem, where full structural alignments are used to find templates, and the quality of the alignment at the interfacial region of the template is used to improve the quality of the prediction (Hosur *et al*., 2011; Zhang *et al*., 2013; Guerler *et al*., 2013; Mirabello and Wallner, 2017). Other methods focus on structurally aligning a subset of residues of the target structures corresponding to the interfacial regions of templates as a more flexible way of finding interaction templates whenever the evolutionary relationship between targets and templates is unclear or non-existent (Günther *et al*., 2007; Gao and Skolnick, 2010a; Tuncbag *et al*., 2011).

However, thus far the full potential and the characteristics of interfaces have not been explored, and it is still unclear whether aligning interfaces, rather than full monomers, represents a real improvement in the search for interaction templates (Sinha *et al*., 2010). In the latest CASP12/CAPRI experiment (2016) the most successful groups were still using templates gathered from alignments of sequences or full structures (Lensink *et al*., 2017). A possible reason for why the full potential of interfaces has not yet been exploited can be found by analyzing how current interface alignment methods are implemented. For example, PRISM (Tuncbag *et al*., 2011), one of the leading methods based on interface alignments, does not restrict its search for templates to only interfacial residues, but also includes neighboring residues that are closer than 6Å to other interfacial residues, even if these are buried in the protein core. Although such an approach helps the alignment procedure by reducing the fragmentation of the interface, it also restricts the alignment to cases where basically the secondary structure elements at the interface level match, and thus it might be less effective at finding evolutionary unrelated, yet compatible, templates.

Another approach to find templates is to rely on using sequence-independent structural alignment programs such as TM-align (Zhang and Skolnick, 2005) and restricting the alignment to only interfacial residues (Kundrotas and Vakser, 2013; Guerler *et al*., 2013). The ability to insert multiple gaps in structural alignments can help to address the issue of fragmentation at the interface. However, depending on the level of fragmentation, TM-align might still fail in retrieving the correct alignment, simply because it is not designed for dealing with heavily fragmented coordinate sets. A further limitation of TM-align is that the sequential order of the residues must be maintained for the algorithm to work correctly, and as such it would not be possible to align interfaces with different chain topologies.

To address the topological issue, iAlign was developed (Gao and Skolnick, 2010a). It is a protein-protein interface comparison method based on an extension of the Kabsch algorithm (Kabsch, 1976), also used in TM-align, that will optionally allow for sequence-order independent comparisons for interfaces. However, it utilizes a definition of protein interface, where the interfacial residues are collected across both the protein chains involved in the interaction. This means that the mutual position of the patches of interfacial residues must be known before any comparison can be performed against a template. Thus, iAlign can only be used if the complete interface is known, e.g. for comparing known interfaces, and not for searching for interfacial residues on a monomer structure.

In this study, we present InterComp, designed to perform sequence-order independent structural comparisons and alignments. Since the algorithm works on a disjointed set of points in space rather than on a sequence of residues, the method is general and can be used for various structural comparisons applications, including interface-surface, interface-interface, and even small molecule ligand comparisons. The main difference in these applications would be a few size-dependent parameters and the statistics, i.e. p-value calculation. Here, we demonstrate a case when we apply InterComp to find protein interfaces on monomer structures (interface-surface). We show that InterComp can pinpoint the interface location on the surface of proteins, even when no templates can be found with the already available sequence-order dependent structural alignment methods (e.g TM-align).

## 2 Methods

### 2.1 Algorithm

The aim of this study is to build software that is capable of comparing molecules by treating them as a set of independent points on a surface in a 3D space. This means that the points have no inherent ordering. This differs from regular structural alignments, where the atoms follow a fixed order according to the protein sequence.

We use a modified version of a stochastic method for molecular structure matching (Kirkpatrick *et al*., 1983; Barakat and Dean, 1991) and simulated annealing to solve an optimization problem that maximizes the structural superposition score of two molecules independently of the chain topology of their covalently bonded network. The objective function is calculated by comparing the *C_α_* distance maps of the two molecules. This simplifies the problem; since distance maps are invariant to rotations and translation there is no need to apply spatial transformations to superimpose molecules. Instead, the optimal matching between two molecules is found by permutating the rows and columns in the distance matrix while maximizing the similarity (see below for details).

### 2.2 Objective function

In this study, the similarity measure for a given trial alignment between two molecules *a* and *b* with length *L_a_* = *N* and *L_b_* = *M* (*N* ≤ *M*), represented as internal distance matrices *D_a_* and *D_b_*, is a variant of the Levitt-Gerstein score (Levitt and Gerstein, 1998) adapted to internal distances:

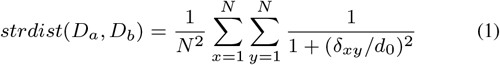

where *N* is number the of residues in *a*, and *d*_0_ is a parameter that monitors the slope of the function (optimized *d*_0_ =0.5Å, see Supplementary information), *δ_xy_* is the absolute element (*x, y*) in the matrix difference between *D_a_* and the first *N* columns and rows of *D_b_* after it has been permutated to form a trial alignment, see diff (*D*_1_, *D*_2_) in Figure 1a. The *M* − *N* residues (for *x* > *N* and *y* > *N*) from *b* are not included in the alignment and are thus excluded from the similarity score.

**Fig. 1.**
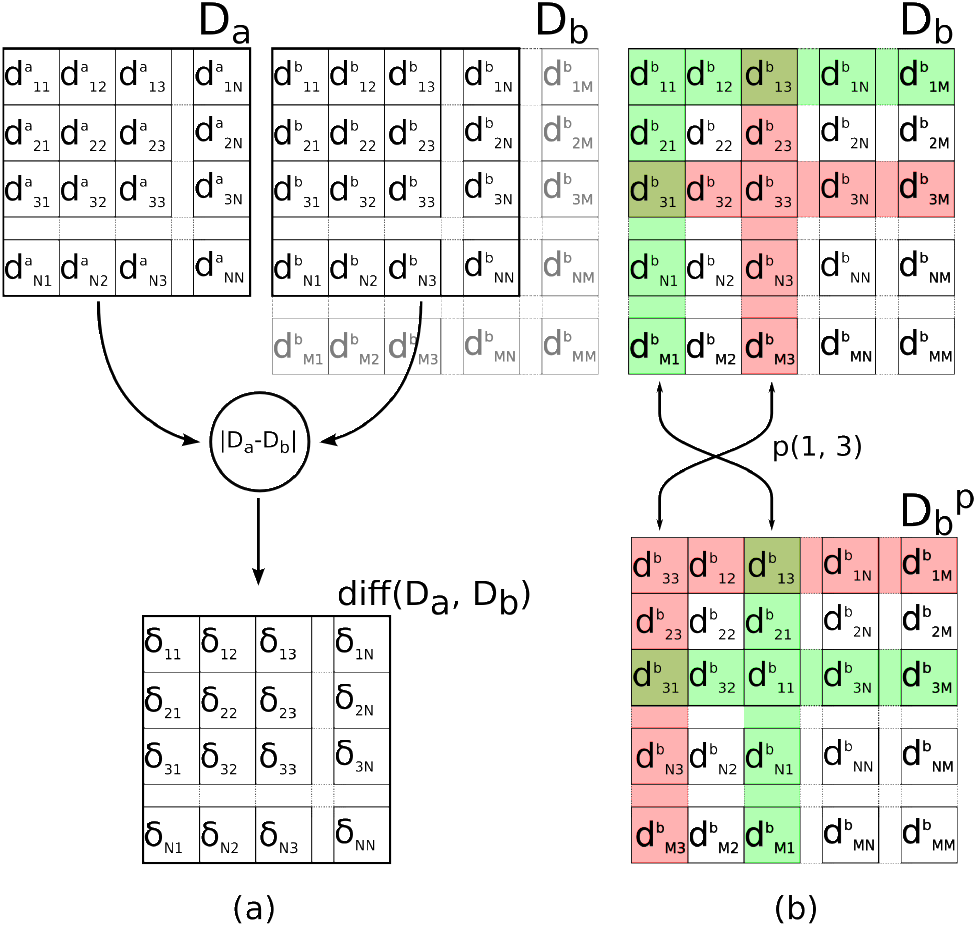
(a): the similarity score in InterComp is calculated by adding up the elements in the matrix of deltas diff (*D_a_, D_b_*). The deltas are the absolute difference, calculated element by element, between the first *N* rows/columns in the distance matrix *D_a_* and the matrix *D_b_*. (b) A trial alignment is obtained by permutating *D_b_* by swapping any two random rows/columns forming the 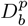

To also consider the chemical compatibility of the molecules (i.e. the similarity of the aligned residues), a second scoring component based on amino acid similarity is used, analogous to the method used for the structural similarity above:

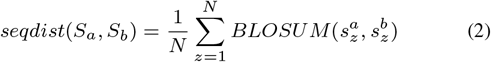

where *N* is the number of residues in *S_a_, S_a_* and *S_b_* are the aligned amino acid residues from targets *a* and *b*, respectively, and the BLOSUM62 substitution matrix (Henikoff and Henikoff, 1992) is used to score the similarity for the matching position.

To represent both the structural and sequence similarity, the two scores are combined using a weighted sum to form the complete objective function in the simulated annealing procedure:

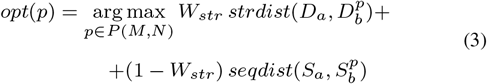

where *W_str_* ∈ [0, 1] is the weight for the structural similarity score, 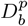 is a permutation of the rows/columns in the distance matrix *D_b_* and 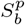 the corresponding amino acids. The default weight of the structural component of the scoring function, *W_str_*, has been optimized to 0.5 by trying different values for *W_str_* in the 0.25 − 1.75 range (see Supplementary information).

When the optimal mapping between two molecules has been found, a structural superposition can be performed by minimizing the RMSD for the mapping and outputting the two structurally aligned molecules in PDB format.

### 2.3 Optimization protocol

The optimization procedure keeps the distance map *D_a_* of the smallest molecule fixed, while trial configurations for the largest map *D_b_* are generated by swapping a random pair of rows/columns. To account for the difference in size, a number of rows/columns in *D_b_*, equal to the size difference, are always ignored when calculating the final score. These are initialized randomly, and then sampled naturally by the swapping of rows/columns in *D_b_*. Figure 1b shows how a trial configuration is generated to obtain 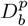 from *D_b_*. In addition, to model mismatches in *D_a_* a proportion of the rows/columns from the smallest molecule, *D_a_* can optionally be ignored (*null correspondences* as defined in (Barakat and Dean, 1991)). However, in our tests this option did not yield better results, most likely because the structural similarity score (Eq: 1) already down-weight residues with large deviations (data not shown).

The simulated annealing procedure is used to find the optimal sorting for *D_b_*. At each iteration, two columns in *D_b_* are randomly swapped to form a trial configuration, the score (Eq:3) is calculated and the configuration accepted if it improves on the current best score. Otherwise, the configuration is accepted with probability: *P* = *exp*(−Δ*score*/*T*), where the Δ*score* is the difference between the trial score and the last accepted score, and *T* is the annealing temperature, which is gradually lowered with the number of iterations.

With regard to the annealing procedure discussed in Barakat and Dean, 1991, the main differences in the current method are the larger number and length of Markov chains used during the search. This is necessary since it is difficult to compare molecules containing many atoms and a large imbalance in the number of atoms between targets. The longer Markov chains allow the state of the system to settle as the temperature is decreased in the annealing procedure. The procedure is said to have converged whenever the acceptance rate, i.e. the frequency at which a new trial configuration is accepted, drops below 0.3%. The acceptance rate is reset at the beginning of each Markov chain, and the minimum size of the Markov chain is set so that the acceptance rate can always drop below the stopping criteria.

### 2.4 Data sets

To test and optimize the method, a data set of hetero- or homo-dimeric protein-protein complexes was constructed. The set is composed of protein chains extracted from a 20% redundancy-reduced version of PDB compiled by PISCES (Wang and Dunbrack, 2003). The reduced PDB contained 2,952 protein chains (July 13, 2016) with resolution 1.6Å and R-factor 0.25 or better. From this set, any protein chain involved in one or more dimer interactions was selected, resulting in 668 protein chains involved in dimer interactions. To avoid including targets with very small interfaces, any monomer with an interface composed of less than 20 residues was removed. In addition, targets with an interface covering more than 50% of their surface were also removed, since these targets were deemed too easy and even a random predictor would score well in our tests. The final set called *T568* consisted of 568 protein monomers. From this, a *monomer shells* set (*S568*) was constructed, consisting of the *Cα* atoms from residues on the surface of the protein chains defined by Residue Solvent Accessibility (RSA) for the side-chain > 15% calculated using Naccess (Hubbard and Thornton, 1993).

In addition, a library of template protein interfaces was also extracted from PDB (May 19, 2016) using the defined biological units to prevent non-native interfaces from crystal packing (Carugo and Argos, 1997). If a biological unit is also a multimer, an interface is defined as residues within 5.0Å (all-atoms) between the two monomers. Each resulting interface is characterized by the monomer from which its residues were extracted and the monomer containing the counterparts in the interaction. For example, if the PDB 1*xyz* contains two chains, *A* and *B*, two separate interfaces will be extracted: 1*xyz_AB*, containing the *Cα* atoms of residues from *A* that were interacting with any residue in *B*, and 1*xyz_BA*, containing the *Cα* atoms of residues from *B* that were interacting with any residue in *A*. The resulting template library, called *I400k*, contained approximately 400,000 interfaces.

Figure 2a shows an example of a protein monomer (PDB id: 3vsv, chain A, colored in purple) where the shell atoms are highlighted as green spheres and the interface atoms (interface id: 3vsv_AD) are highlighted in red. It is important to note that all the interface atoms are also shell atoms. A stylized version of the same concept is shown in Figure 2b. In this work we use InterComp to align the interface to shells (Figure 2c), and compare InterComp to using TM-align to align the interface to a monomer (Figure 2d) and monomer to monomer (Figure 2e).

**Fig. 2.**
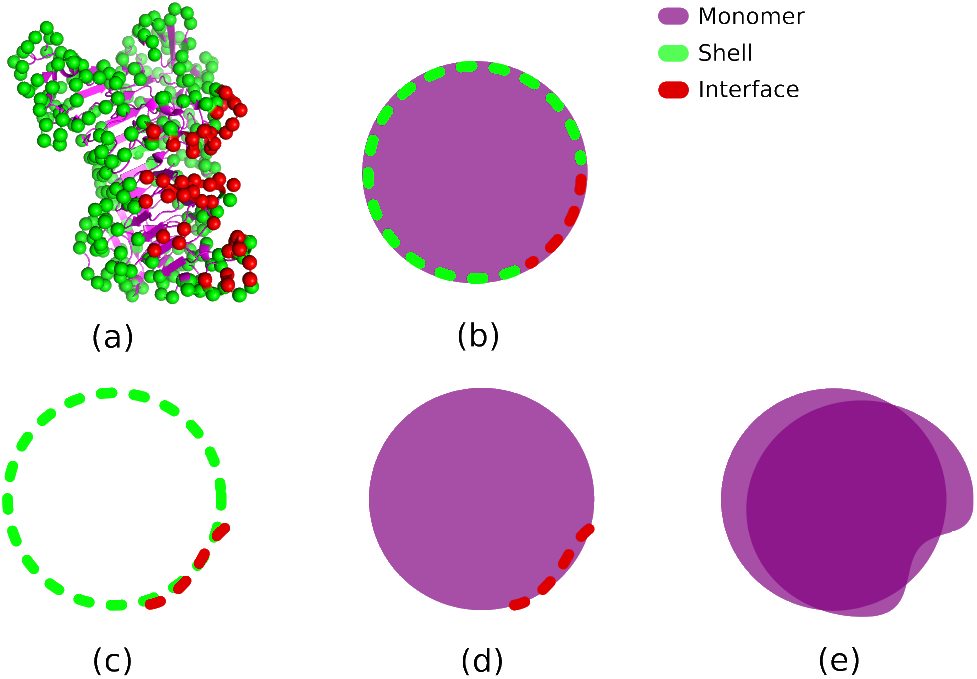
(a): Protein monomer 3vsv, chain A (purple). The shell atoms are *Cα* atoms from residues whose RSA is over 15% (green). The interface atoms are a subset of the shell atoms whose residues are also closer than 5Å to any residue in another protein chain (red). (b): Stylized version of a protein monomer where the shell and interface residues are represented with a dashed contour line. (c): Example of an interface-shell alignment, where an interface and a shell extracted from two monomers are aligned. This is the case with all InterComp alignments in this work. (d): Example of an interface-monomer alignment, where the interface atoms are aligned against all *Cα* atoms in a monomer. (e): Example of a monomer-monomer alignment, where all *Cα* atoms from two protein monomers are aligned. The alignments shown in d and e are performed with TM-align in this work.

In order to estimate p-values for the structural score (Eq:1), a set of approximately 2 million random alignments between target shells and biological interfaces was constructed. This set was constructed by aligning 1,790 monomer shells previously described in Gao and Skolnick, 2010a against the library of biological interfaces (I400k) using InterComp to calculate the structural score. To avoid including non-random hits, pairs that showed a significant similarity by TM-align (TM-score<0.35) were filtered out. It is important to note that this filtering will only remove any high scoring interface-shell alignment that can be found by TM-align (sequence-order dependent). It is still possible that the set of random alignments may include non-random high scoring matches, interfaces that are only found when compared in a sequence-order independent manner using InterComp. Still, these examples will be few compared to the whole random set and should not influence the p-value calculation too much.

Since the structural score depends heavily on the size of the interface and shell, the random structural scores were fitted to an extreme value distribution for different interface and shell size bins. In Fig. 3 the random distribution for different size bins along with the fitted distributions are shown. We used the fit to calculate the p-values for each structural score. By analyzing the first row of plots in Fig. 3, it is clear that whenever interfaces contain less than 10 residues, independently of the size of the shell, random scores tend to be high. This will make it hard to find significant hits for small interfaces, which makes sense, since smaller interfaces can easily align with most targets (e.g. single helix to helix alignment).

**Fig. 3.**
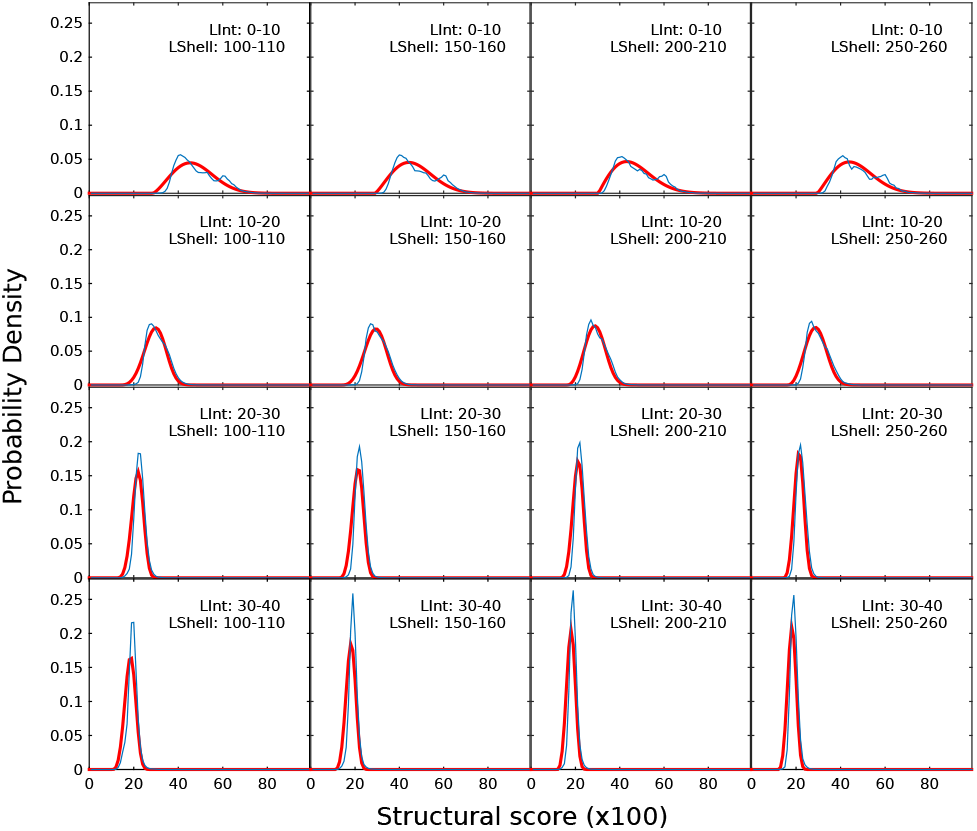
Probability density distributions of InterComp scores from random alignments for different interface (rows) and shell (column) sizes. The empirical probability density is shown in blue and the fitted extreme value distribution in red.

Since our tests consist of multiple comparisons against many templates (400k), using the p-value alone to decide whether the null hypothesis can be rejected or not would be impossible without performing the necessary multiple testing correction. In our tests, we thus adopt the False Discovery Rate (FDR) controlling procedure (Benjamini and Hochberg, 1995) and derive an FDR adjusted p-value (q-value) for each p-value (Yekutieli and Benjamini, 1999). The q-values are then used to decide if a given InterComp score is significant.

To test the hypothesis that InterComp is more sensitive than a sequence-order dependent algorithm, two additional interface sets of varying difficulty were constructed. The first “medium” set was built by removing all template hits that would also be matched by TM-align (TM-score>0.5) for a given target from the T568 set, using *Monomer-Interface* alignments, see Fig. 2d. The second “hard” set was built by removing all template hits that would also be matched by TM-align (TM-score>0.5) for a given target from the T568 set, using *Monomer-Monomer* alignments, see Fig. 2e.

In Fig. 4 a schematic representation of all three subsets of template interfaces used to test InterComp is shown. It is important to note that while the I400k is the same for every target in the T568 set, the “medium” and “hard” sets will vary from target to target, depending on the structural similarity found using interface-monomer and monomer-monomer alignment with TM-align.

**Fig. 4.**
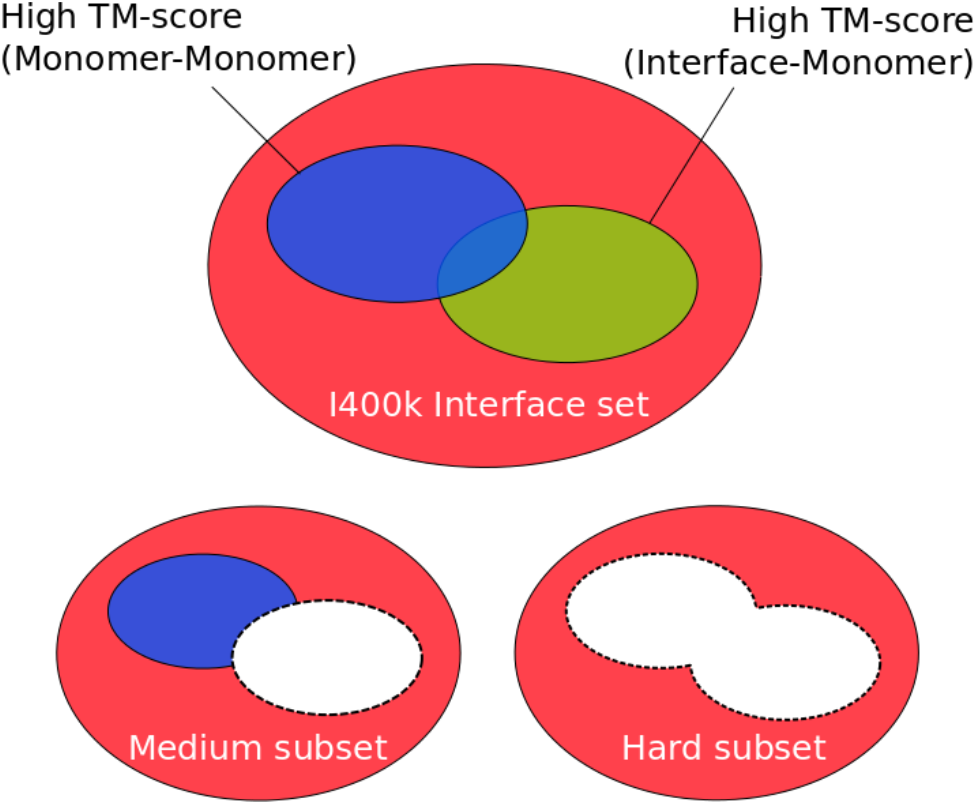
Schematic overview of the different template interface sets that were used to test InterComp. The full I400k interface set, also called the “easy” set, is shown in red. The “medium” subset of I400k for a given target is obtained by removing any template interface that will align with the TM-align, TM-score of the Interface-Monomer alignment ≥ 0.5. The “hard” subset of I400k is obtained by removing from the “medium” subset any template interface whose parent monomer is in the same fold as the target monomer, TM-score of the Monomer-Monomer alignment ≥ 0.5.

### 2.5 Performance measures

To assess the performance of InterComp, the Positive Predicted Value (PPV) of interfacial residues is used:

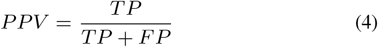

where *TP* is the number of *True Positives*, i.e. the number of residues correctly predicted as part of an interface, and *FP* is the number of residues incorrectly predicted as part of an interface. The sum of *TP* and *FP* is also the total number of residues predicted as part of an interface.

## 3 Results and Discussion

### 3.1 Benchmark: Aligning interfaces to surfaces

InterComp was applied to the problem of predicting the interface residues on the surface of protein chains involved in a dimeric interaction. By restricting the search to the surface of proteins (shells) rather than to full monomers, the complexity of the search was reduced. This was necessary since the time required for the algorithm to converge is related to the length of the Markov chains which increases quadratically with the size of the largest target. The accuracy should not suffer from this choice, since interfaces are expected to be on the surface of proteins.

We hypothesize that top-ranking interface templates by q-value (see Methods) should correspond to residues involved in interactions with partner proteins. The location of the interface on each monomer was found by searching for areas on the surface that are compatible in shape and chemical composition to interfaces in the library, as illustrated in Fig. 2c.

In detail, InterComp was used to align each monomer target shell from the *S568* set against the *I400k* interface library (“easy” test). The top 1 and top 10 best matches by q-value are selected and the performance is assessed by the fraction of correctly predicted interface residues from the target shell, PPV (Eq:4), see Fig. 5 and Fig. 6 right panel. To account for the fact that the “easy” test database contains several simple cases that any structural alignment method would find. The test was made more difficult by removing any template that would be found using TM-align, a commonly used sequence-order dependent structural alignment method. Two difficulty levels were tested: for the first (“medium” test) any template with an interface-monomer TM-score>0.5 was removed (Fig. 5 and Fig. 6 middle panel); for the second (“hard” test) any interface whose parent monomers had a TM-score>0.5 with the target was removed, effectively removing any templates that overall were structurally similar to the target (Fig. 5 and Fig. 6 right panel). In this way, it is possible to assess if InterComp is capable of finding correct template interfaces even when sequence-order dependent structural alignment methods like TM-align fail at various levels.

**Fig. 5.**
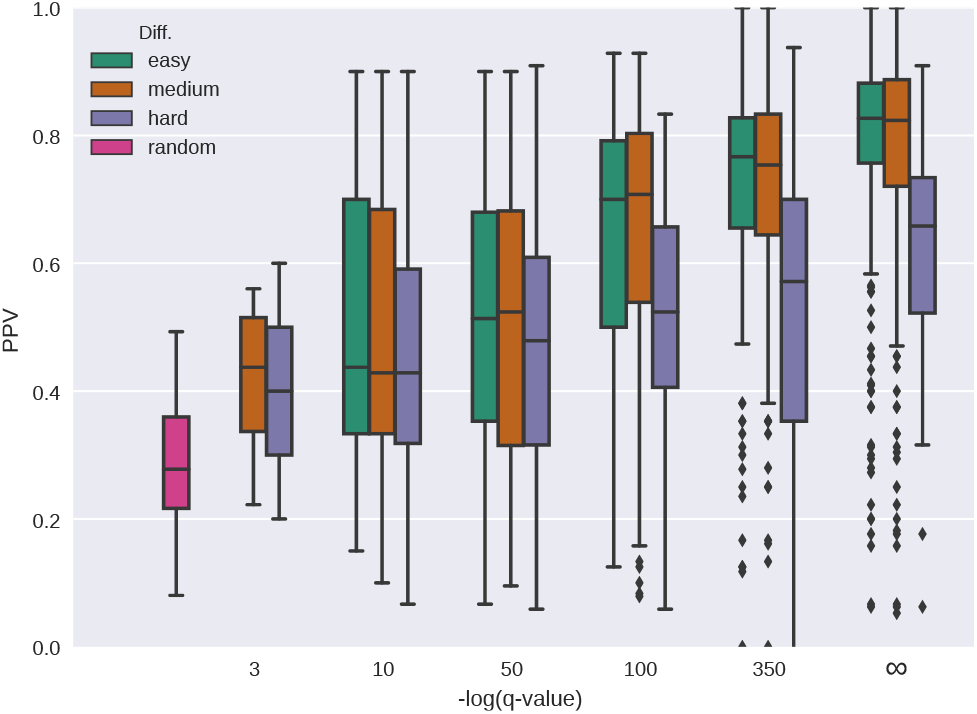
Distribution of fraction of correctly predicted interface residues (PPV) in relation to the negative logarithm of the q-value calculated from the top 10 InterComp hits using the “easy” (green), “medium” (orange), and ‘hard” (purple) interface sets. The “random” box corresponds to the fraction of interfacial residues on each target shell and estimates the performance of a predictor that would pick random shell residues to be part of the interface. The infinity (∞) sign on the x-axis corresponds to q-value=0.

**Fig. 6.**
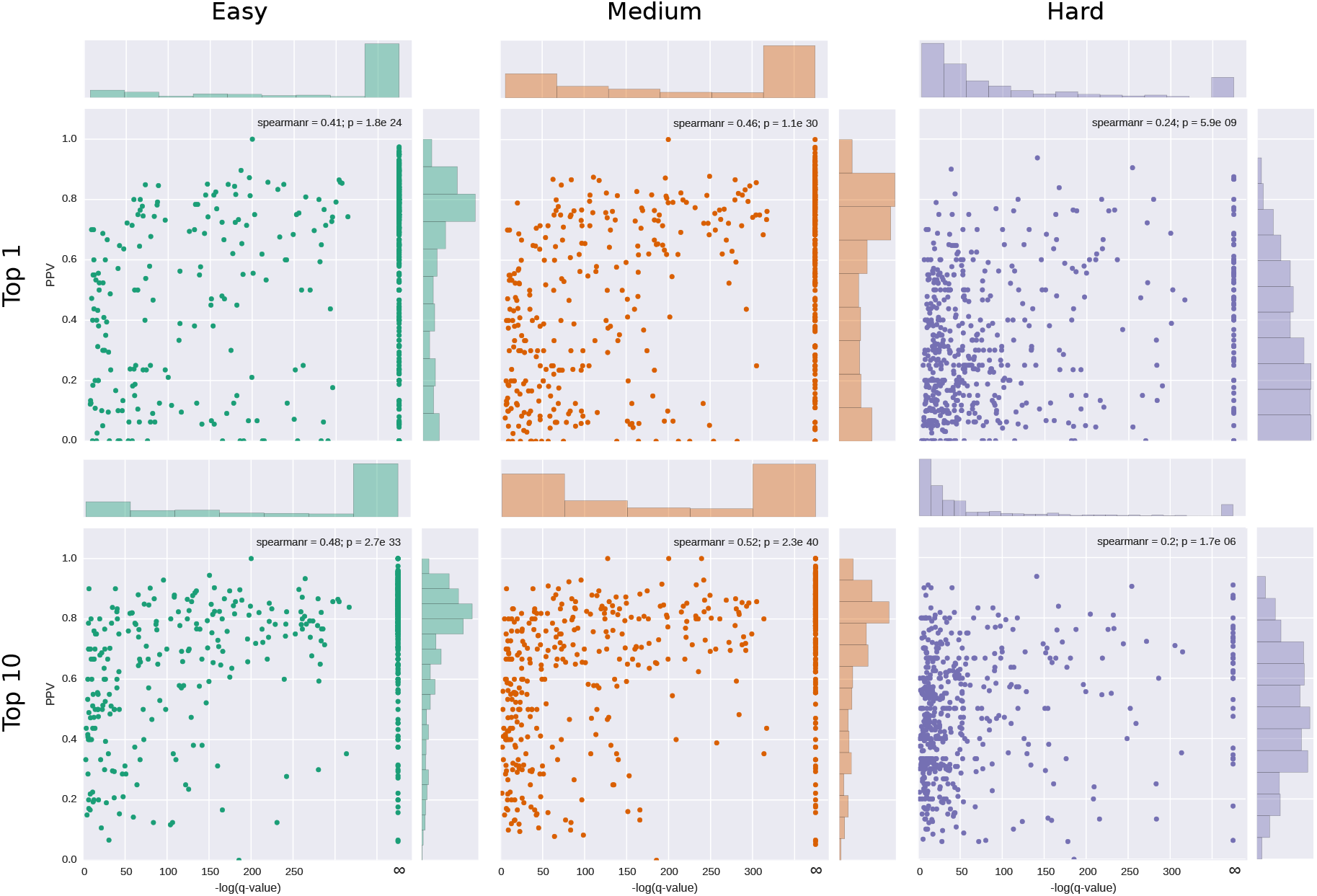
Scatter plots of the PPV in relation to the negative logarithm of the q-value calculated from the InterComp score for the Top 1 and Top 10 hits for each target shell using the “easy” (green), “medium” (orange), and “hard” (purple) interface sets. The infinity (1) sign on the x-axis corresponds to q-value=0.

There is a clear correlation between the structural score from InterComp and the PPV for all three tests (Fig. 5 and 6). Although the correlation between InterComp score and PPV for the “hard” test, is less pronounced (Spearman’s rank correlation 0.20-0.24), InterComp could still find at least the location of an interface with PPV≥0.5 for more than half (287/568) of targets for the top 10 hits and for 24% (139/568) of the targets for the top 1. Moreover, the chances of finding the right interface increases as the q-value derived from the InterComp improves (Fig. 5, purple boxes). For the “medium” test, InterComp could correctly identify the correct interface for 70% (401/568) of targets for the top 10, and 60% (341/568) for the top 1. On the full I400k template set (excluding self-hits, “easy” test) InterComp founds at least one interface for 80% of targets (450/568) and 66% (373/568) for the top 10 and top 1 hits, respectively. In all cases, including the “hard” set the number of correctly identified interfaces is significantly better than would be expected by a random predictor.

### 3.2 Alternative Interfaces

In some cases, even at very significant q-values the PPV can be low or close to zero (Fig. 6). The reason for this could of course be that InterComp predicts completely wrong interfaces in all of these cases. However, a more likely explanation is some of these very significant hits are alternative, yet unknown interfaces or interfaces not included among the test interfaces for the target. An example of a potential correctly predicted interface that is classified as incorrect in the benchmark is shown in Fig. 7a. The top 1 interface predicted by InterComp (q-value=0) for target 4b1y (medium difficulty) is incorrect, since it does not match the interface in its biological unit. However, the monomer from which the template interface was extracted superimposes almost perfectly with the target, as shown in Fig. 7b. This highlights a problem with multiple correct interfaces, and although some can be identified, not all of them are included in the relevant biological unit.

**Fig. 7.**
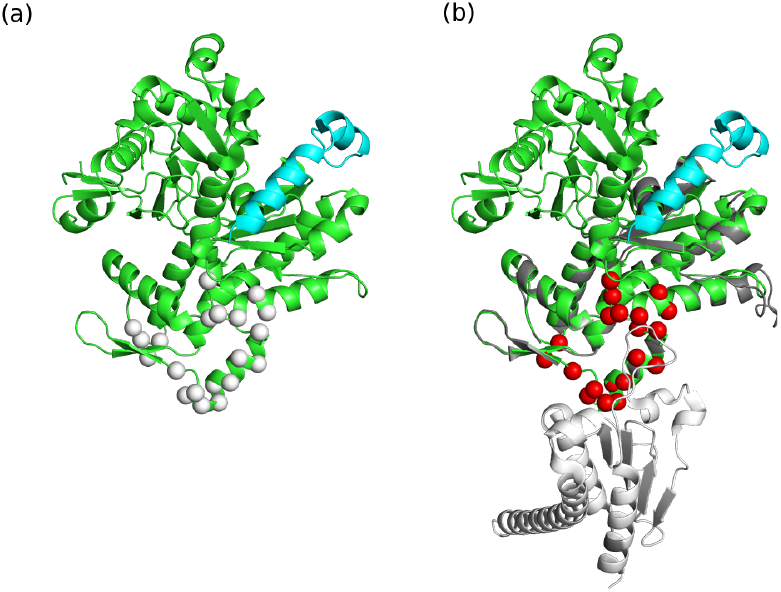
An example of a significant hit that is not part of the target set. (a) The protein Phactr1 RPEL-3 (4b1y) from organism Oryctolagus cuniculus bound to G-actin (cyan). The target interface is the binding site for G-actin (short peptide shown in cyan), but InterComp identified another high-confidence interface with template 2p9l1_BF (white spheres). However, the template 2p9l (Arp2/3 complex from organism Bos taurus) superimpose almost perfectly to the target (green/grey chains aligned on right side, RMSD: 0.65), with the interface 2p9l1_BF highlighted in red and the relevant partner for the template interaction shown in white. In this case, it is likely that the inferface pinpointed by InterComp is actually correct, even though it was not part of the target set.

### 3.3 Successful Examples

A few examples of successful cases from the “hard” tests are shown in Fig. 8. For all of these examples, InterComp is able to find with high accuracy (PPV>0.5) the location of the target interface. This would not have been possible using TM-align, since the TM-score is less than 0.5 for aligning the template interface to the target (0.23-0.35) and for aligning the template interface parent monomer to the target (0.26-0.42). It is easy to spot the location of some false positives, where the predicted interface residues (the red spheres) are quite far away from the interaction surface, e.g. on the rear side of 1t92A in Fig. 8a and 3lagA in Fig. 8b. In these cases, a simple clustering technique could potentially be used to filter out spurious positives that are unlikely to be part of the interface. But that is not the focus of this study.

**Fig. 8.**
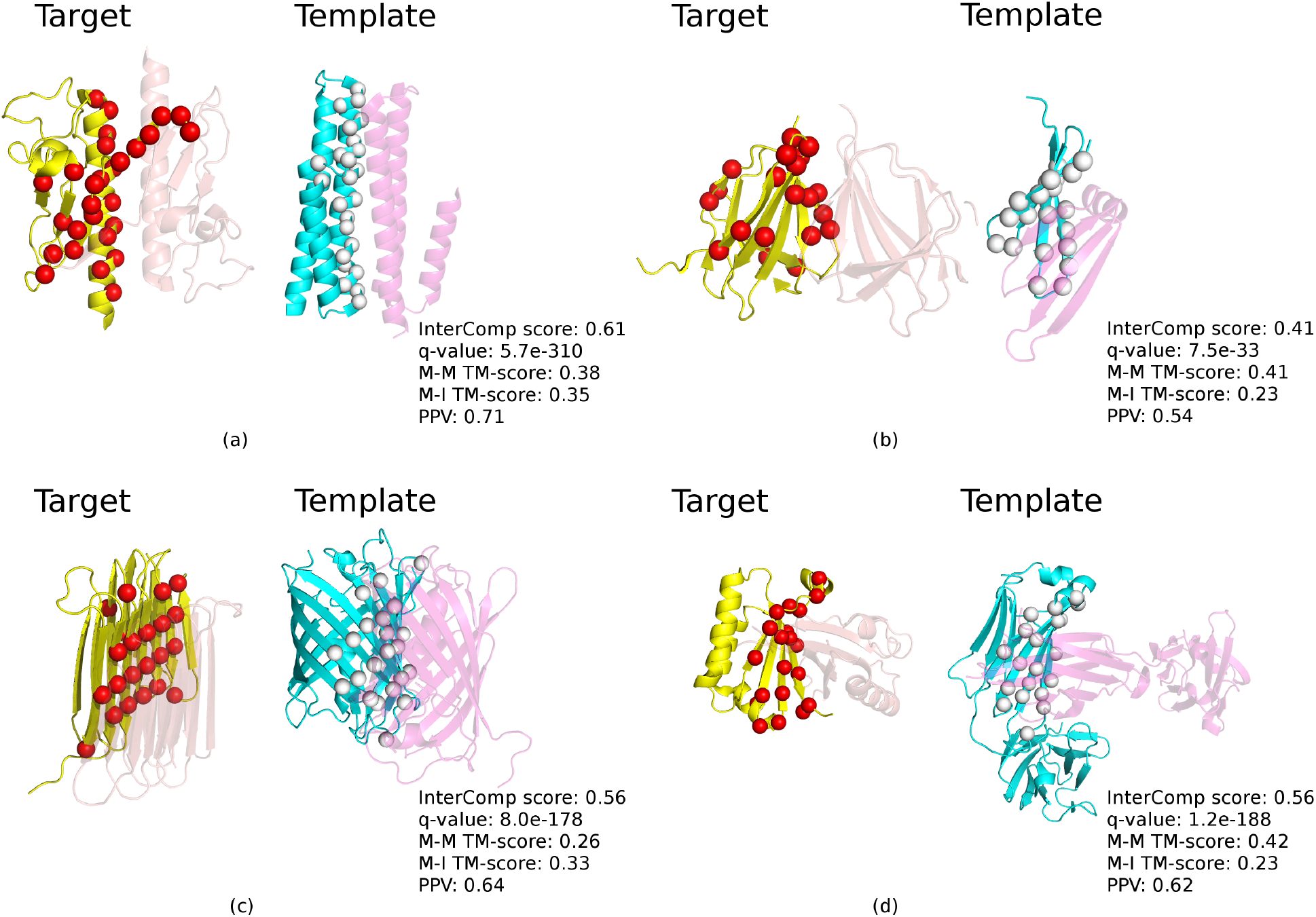
Successful predictions from the “hard” category with the corresponding InterComp score, q-value, M-M TM-score (Monomer-Monomer TM-score), M-I TM-score (Monomer-Interface TM-score) and PPV. The target chains are shown in yellow with the predicted interfacial residues highlighted by red spheres, template chains are shown in cyan with the aligned interface residues highlighted by white spheres. The interacting target partners are showed in transparent orange and template partners are showed in transparent magenta. (a) Target 1t92A, Pili subunit structure of N-terminal truncated pseudopilin PulG from Klebsiella oxytoca. Template 4uxz1_BC, crystal Structure of a Membrane Diacylglycerol Kinase from organism Escherichia coli. (b) Target 3lagA, Double-stranded beta-helix, crystal structure of a functionally unknown protein RPA4178 from Rhodopseudomonas palustris. Template 3kvp1_AD, Beta-propeller-like, crystal structure of uncharacterized protein ymzC precursor from Bacillus subtilis. (c) Target 4dt5A, Single-stranded left-handed beta-helix, crystal structure of antifreeze protein from Rhagium inquisitor. Template 3cgl1_DC, GFP-like protein dsFP483, cyan fluorescent protein from organism Discosoma striata. (d)Target 5b08A, Alpha-beta plaits, polyketide cyclase OAC from organism Cannabis sativa. Template 3oay2_LM Immunoglobulin-like beta-sandwich, HIV glycan shield from Homo sapiens.

### 3.4 Computational cost

To give an idea of the computational cost of running InterComp, we timed the running times using a few typical sizes for target shells against the full I400k set of template interfaces using a 28 core 2.6 GHz computer with 128GB RAM Linux node. The median number of residues on the surface of a monomer (shell size) for set S568 is 129. This corresponds to a monomer of about 200 residues (e.g. target 2zcmA, 3cxnA) with a runtime of approximately 120 minutes on 28 cores (56 core hours in total). A run on a target shell twice as large (250 residues, and 400 residues in total in the monomer) will need approximately 13 hours to be completed (364 core hours in total). Overall, a full test on all 568 targets in the S568 set against I400k takes approximately 100,000 core hours, i.e. 176 core hours per target on average.

## 4 Conclusions

We have presented InterComp, a topology and sequence-order independent structural alignment method. We have shown that InterComp is capable of performing protein surface to interface alignment and can be used to pinpoint potential interaction points on the surface of proteins, even when regular structural alignment methods that are dependent on the sequence-order fail. The fact that InterComp can align monomer structures to one side of a complete interface is extremely useful, and should leverage the use of structural information in protein-protein docking by providing novel templates with similar interfaces but no overall structural similarities. However, the interface-surface alignment case demonstrated here is only one of many potential use-cases for InterComp. For instance, we are currently recalculating the statistics to apply the method to interface-interface, and small molecule comparisons, which will enable clustering of interfaces and improvements to virtual screening of small molecules.

## Acknowledgement

This work was supported by Swedish Research Council grants 2012-5270, 2016-05369, The Swedish e-Science Research Center, and the Foundation Blanceflor Boncompagni Ludovisi, née Bildt. The computations were performed on resources provided by the Swedish National Infrastructure for Computing (SNIC) at the National Supercomputer Centre (NSC) in Linköping. We also thank Isak Johansson-Åkhe for helpful discussions.

## Supplementary Information for “InterComp: sequence-order independent structural matching applied to protein interface alignment”

### S1 Parameter optimization

InterComp has three parameters that have a significant impact on the search for the optimal mapping between two molecules. The first parameter is *d*_0_, which is a parameter of the Levitt-Gerstein score (see Eq. 1), the second parameter is the weight of the structural component of the scoring function in relation to the weight of the chemical component, *W_str_* (see Eq. 3), and the third parameter is the percentage of null correspondences, i.e. the number of rows/columns that can be ignored during the annealing procedure in the distance matrix of the smallest of the two molecules. It should be noted that the weight on the sequence is 1 − *W_str_*, which means that increasing the structural score will in effect decrease the relative contribution from the sequence similarity. The default values for *d*_0_ = 0.5 and *W_str_* = 0.5 were found by a grid search trying all combinations of *d*_0_ ∈ [0.25, 1.75] in 0.25 steps and *W_str_* ∈ [0.50, 0.75, 1.0] maximizing the number of target monomers whose interfaces were correctly (PPV>0.5) found within the top 10 identified templates (Fig. S1). In addition, the combination *d*_0_ = 0.5 and *W_str_* = 0.25 was also tested (the single low-performing point in the “hard” category in Fig. S1), but considering the bad performance for the “hard” category and the high computational cost for obtaining additional points (35,000 core hours per point) no more combinations involving *W_str_* = 0.25 were tried. The default values *d*_0_ = 0.5 and *W_str_* = 0.5 were selected to allow for better results in most cases (mainly on the “easy” and “medium” sets). Still, a higher weight on the structural component of the score (*W_str_* ≥ 0.75) or a higher *d*_0_ could yield slightly better results in the “hard” category. For the optimized *d*_0_ and *W_str_* allowing for 10% null correspondences, did not improve the performance (data not shown). Thus, the percentage of null correspondences was set to 0.

**Figure S1:**
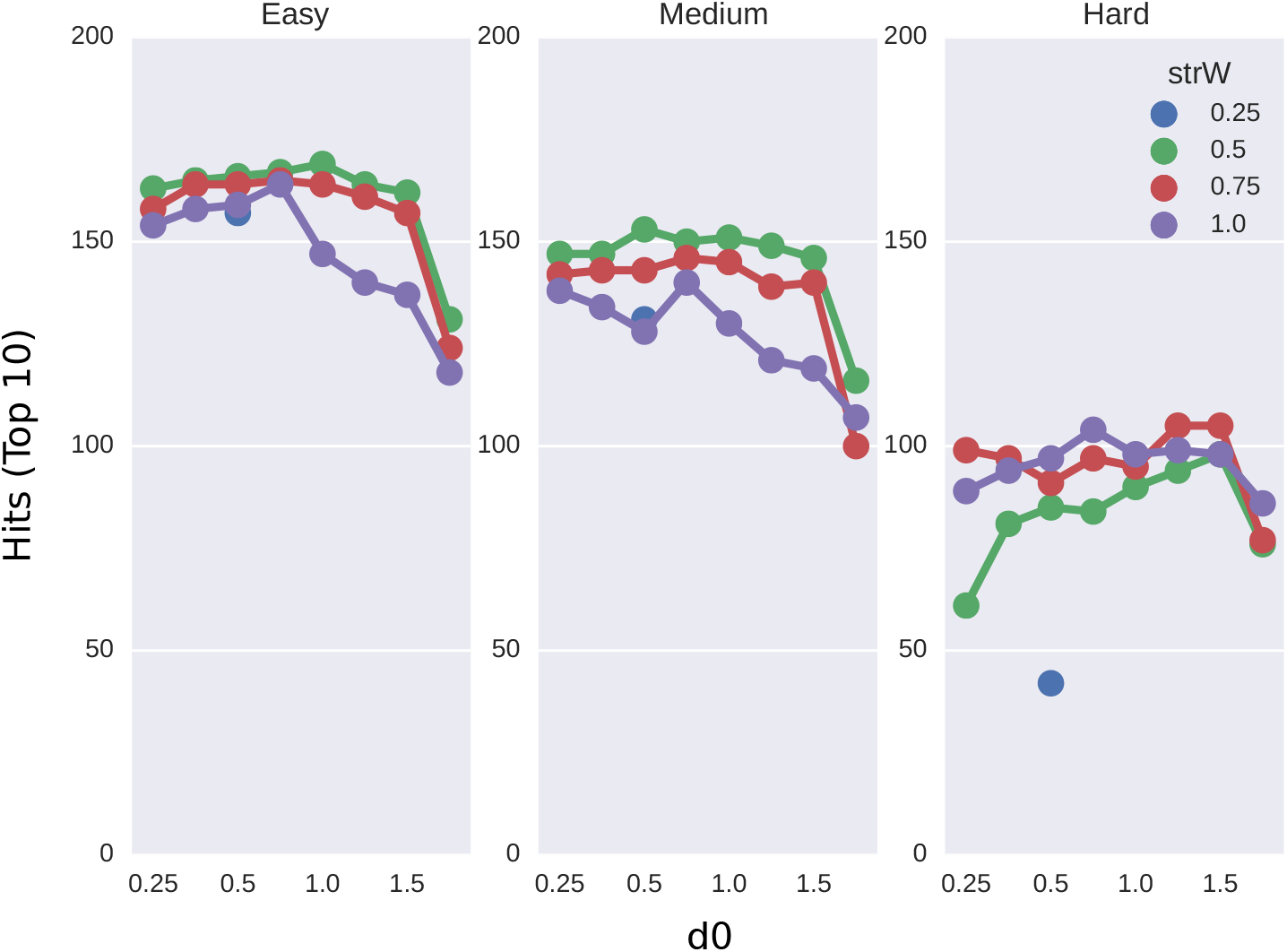
Number of targets for which the interface could be correctly detected at various parameter settings. This test was run on a subset of 200 targets randomly selected from set S568. The selected default parameters are *d*_0_ = 0.5, and *W_str_* = 0.5.

